# HLA-DQ monomer-based enrichment of alloreactive CD4 T cells

**DOI:** 10.1101/2025.05.30.656890

**Authors:** Zhuldyz Zhanzak, Davide Cina, Xueqiong Zhang, Annette Hadley, Haydn T. Kissick, Christian P. Larsen

## Abstract

Donor-specific T cell responses, particularly against human leukocyte antigen (HLA)-DQ antigens, are critical in transplant immunology as they influence graft survival and rejection. This study investigated HLA-DQ-specific CD4 T cell responses in naive individuals using HLA-DQ monomers, representing common donor HLA-DQ antigens (DQ2.5, DQ5, DQ6, DQ8), as defined alloantigen sources to enrich and detect HLA-DQ-specific T cells. Using a repeated stimulation protocol with HLA-DQ monomers, we achieved an enrichment of HLA-DQ-specific CD4 T cells, evidenced by proliferation and activation marker upregulation. Additionally, *in silico* epitope prediction identified 15-mer HLA-DQ peptides capable of binding to self-HLA-DRB1 alleles. TCR sequencing demonstrated an enriched, oligoclonal repertoire of HLA-DQ-specific CD4 T cells post-stimulation, with specific TCR clonotypes expanding in response to HLA-DQ monomer stimulation. Together, our study presents an *in vitro* method for enriching HLA-DQ-specific CD4 T cells using HLA-DQ monomers as an alloantigen source, thereby improving the detection of alloreactive, donor-specific T cells in naive individuals with the potential to mediate immune responses, including graft rejection and donor-specific antibody (DSA) production.

## 1. Introduction

Immunosuppression targeting the T cell alloimmune responses has significantly improved early renal allograft survival (1). Despite the ongoing optimization of T cell-directed immunosuppressive protocols over time, episodes of acute and chronic rejection continue to contribute substantially to allograft injury. Specifically, the evolution of donor-specific antibodies (DSA) directed at human leukocyte antigens (HLA) is a sentinel and irreversible event in the alloimmune response that often leads to progressive allograft dysfunction and ultimately graft loss (2,3). For reasons that remain incompletely elucidated, DSA against HLA-DQ molecules are the most common type of alloantibody after transplantation and correlate most closely with poor graft outcomes (4–7). It is therefore important to characterize the stepwise development of HLA-DQ alloantibodies to better understand immune rejection in transplantation.

Engagement of a peptide-major histocompatibility complex (pMHC) by an antigen-specific CD4 T cell lies at the origin of the antibody response. The task of comprehensively determining HLA-DQ antigenic peptides and identifying their corresponding T cells *in vivo* or *ex vivo* remains a significant challenge for several reasons (8). First, the vastness of the human T cell repertoire, combined with a relatively low frequency of naive T cells specific for any given pMHC, is the principal hurdle to overcome in addressing this problem (9). Second, selecting these rare antigen-specific T cell clones via experimental approaches is rendered more challenging by the relatively weak affinity of the pMHC-TCR interaction (10). Last, the inherently polymorphic nature of HLA both as an antigen and as the context within which antigenic peptides are presented, as well as the polyspecificity of both individual pMHC and T cell receptors (TCR), compound the complexity of the problem (11).

Several empirical and computational approaches have been devised to address this problem largely in the context of infectious disease and cancer. Tetramers and multimers are engineered multivalent MHC molecules designed to present selected peptides to antigen-specific T cells, thereby capturing specific pMHC-TCR interactions. Traditionally, these are conjugated to probes that tag the target cells which are then sorted via flow or mass cytometry (12). Such an approach is limited in throughput by the number of fluorophores or isotope tags that can be multiplexed and signal strength. This has been circumvented by tagging tetramers or multimers with DNA barcodes which can then link a pMHC multimer to a specific TCR by cell sorting and sequencing (13,14).

The addition of microfluidics and droplet-based single-cell sequencing to this approach further optimized the detection of individual rare T cell clones (15). While this approach vastly improves the number of simultaneous pMHC-TCR interactions that can be explored as well as the detection of rare or non-expanded clones in the naïve repertoire, its scalability is hampered by the need to identify and synthesize putative antigenic peptides and MHC multimers. Furthermore, this approach does not capture pMHC-TCR interaction in its endogenous context and appears better suited for studying the interaction between class I MHC and CD8 T cells (12). This issue is of note in the field of transplantation, where the indirect alloimmune response and class II MHC presentation are thought to be a greater contributor to allograft injury long-term (16–18).

Given the marked complexity of experimental methods for determining pMHC-TCR specificity, considerable efforts have been made to develop computational methods for predicting this interaction. These approaches commonly use tetramers (19,20) or dextramers (21) with known antigenic peptides along with structural biology and TCR sequencing to generate training data for an algorithm that identifies TCRs with shared specificity. Such algorithms are then able to not only cluster MHC restricted-antigen specific TCRs in naïve and enriched populations, but also predict TCR sequences directed to a given pMHC complex. While these *in silico* approaches show great promise to increase the throughput of discovery in this space, they still suffer from the inherent limitations of non-native pMHC presentation and have not been tested against antigens as complex and diverse as human HLA.

In this study, we aim to address this challenge by utilizing HLA-DQ monomers to identify and enrich HLA-DQ-specific CD4 T cells from peripheral blood mononuclear cells (PBMC) of naive individuals. We employed a combination of *in vitro* stimulation, epitope prediction, and functional assays to enhance the sensitivity of detecting these T cells. By expanding CD4 T cell lines and mapping TCR repertoires, we sought to elucidate the specific T cell epitopes and clonotypes associated with HLA-DQ recognition. Our approach leverages computational tools and experimental validation to provide a comprehensive understanding of HLA-DQ-specific T cell responses in naive individuals with the view of developing better immune monitoring tools and targeted immune-therapeutics.

## 2. Materials and Methods

### 2.1. Human samples and HLA-typing

Untreated human samples were obtained from Leukocyte Reduction System during the leukapheresis process (StemCell, 200-0093). Leukocytes were further processed using the EasySep Human CD4 T Cell Isolation Kits and EasySep™ magnet (StemCell,19052). Cells were used fresh or cryopreserved in 90% FBS with 10% dimethyl sulfoxide (DMSO), and eventually stored at -150°C.

### 2.2. Flow cytometry

Cells were thawed, counted and stained in Dulbecco’s phosphate-buffered saline (DPBS) + 2% FBS with LIVE/DEAD Fixable Blue Dead Cell Stain Kit (1:1000, Life Technologies) and surface-stain antibodies at 0.5 tests per 1e6 cells in brilliant stain buffer (BD Bioscience, 563794). All data were acquired the same day on FACSymphony A5 with FACSDiva v.8.0.1 (BD Biosciences) and analyzed using FlowJo software (v.10.6.1).

### 2.3. HLA-DQ-specific T cell expansion

Cells from three different Leukocyte Reduction System cone samples were used for *in vitro* T cell expansion and labelled with a Cell Trace Violet Proliferation kit (1:1000, ThermoFisher, C34571) at the beginning of T cell expansion. CD4 T cells were co-cultured with irradiated PBMCs (30Gy) at a 1:1 ratio in serum-free AIM-V medium (ThermoFisher, 12055091) in the presence of rIL-2 (Peprotech, 50 IU/ml), rIL-7 (Peprotech, 25 ng/ml). Cells were stimulated with HLA-DQ monomers- HLA-DQB1*02:01/A1*05:01, HLA-DQB1*03:02/A1*03:01, HLA- DQB1*05:01/A1*01:01, HLA-DQB1*06:02/A1*01:02 (NIH Tetramer Core, 1mg/ml). Cytokines, antigens, and fresh media were supplemented whenever cells were split during the expansion period.

### 2.4. Identification of HLA-DQ epitopes

Potential CD4 T cell epitopes derived from HLA-DQ proteins (HLA-DQB1*02:01/A1*05:01, HLA- DQB1*03:02/A1*03:01, HLA-DQB1*05:01/A1*01:01, HLA-DQB1*06:02/A1*01:02) were predicted using epitopeR package. Presenting elements were used as HLA-DRB1*01:01, DRB1*16:02, DRB1*15:01, DRB1*11:01, DRB1*08:03, DRB1*04:01. 15-mer predicted peptides (**Supplemental Table 2**) were synthesized as crude material and ultimately resuspended in DMSO (Genscript).

### 2.5. Epitope mapping by ELISpot

Expanded and rested cells were plated in an IFN-γ ELISpot (1 × 10^5^ cells per well, Mabtech, ELISpot Pro: Human IFN-γ (HRP), 3420-2HPW-2 and stimulated with peptides for 48 hours (10 μg/ml per peptide). Following the stimulation, the manufacturer protocol was followed step-by-step. After the spot formation, the plate was inspected, and spots were counted in an ELISpot reader.

### 2.6. TCRαβ sequencing

Genomic DNA was isolated from the samples using DNeasy Blood and Tissue Kit (Qiagen, 60504). It was further used for TCRαβ sequencing (immunoSEQ assay; Adaptive Biotechnologies). Amplicons were sequenced using the Illumina NextSeq platform. Raw sequence data were filtered on the basis of TCRβ V, D, and J gene definitions provided by the IMGT database (22).

### 2.7. Quantification and Statistical Analysis

Statistical analysis was done using GraphPad Prism software (Version 10.0.2). The unpaired Mann–Whitney test was used when appropriate and as indicated in each figure legend. p< 0.05 was considered significant. All statistical tests were described in figure legends.

## 3. Results

### 3.1. HLA-DQ monomer-driven expansion enhances the detection of HLA-DQ-specific CD4 T cells

To investigate HLA-DQ-specific CD4 T cell responses in naive individuals, we collected PBMCs from three subjects and performed high-resolution HLA typing to identify each subject’s unique self-HLA alleles (**Supplemental Table 1**). As substantial evidence indicates that HLA-DQ2.5, DQ5, DQ6, and DQ8 are commonly targeted by HLA-DQ DSA, we selected four HLA-DQ monomers representing these prevalent donor HLA-DQ antigens from the NIH Tetramer Core (23,24). These monomers were used to stimulate PBMCs as an alloantigen source, providing a controlled, defined stimulus to assess the specificity and functional response of CD4 T cells. To minimize self-reactivity, we excluded any HLA-DQ monomers that matched the self-HLA alleles of each subject from the stimulation cocktail. CD4 T cells labeled with CellTrace Violet (CTV) were then co-cultured with irradiated PBMCs, which had been pulsed with the selected HLA-DQ monomers and stimulated every nine days across six rounds of stimulation to enrich for HLA-DQ-specific CD4 T cells (**Figure 1A**). This prolonged and stepwise stimulation protocol was designed to progressively increase the precursor frequency of HLA-DQ-specific CD4 T cells, facilitating their identification and further analysis (25). After this expansion protocol, we assessed CD4 T cell proliferation and activation in response to the HLA-DQ monomers. Proliferation was indicated by dilution of the CTV dye, while upregulation of the activation marker CD25 served as an additional indicator of antigen-specific activation (**Figure 1B**). CD4 T cells stimulated with HLA-DQ monomers showed a proliferation relative to unstimulated CD4 T cells, and the addition of an anti-HLA-DR antibody effectively inhibited proliferation (**Figure 1B**).

**Figure 1.**
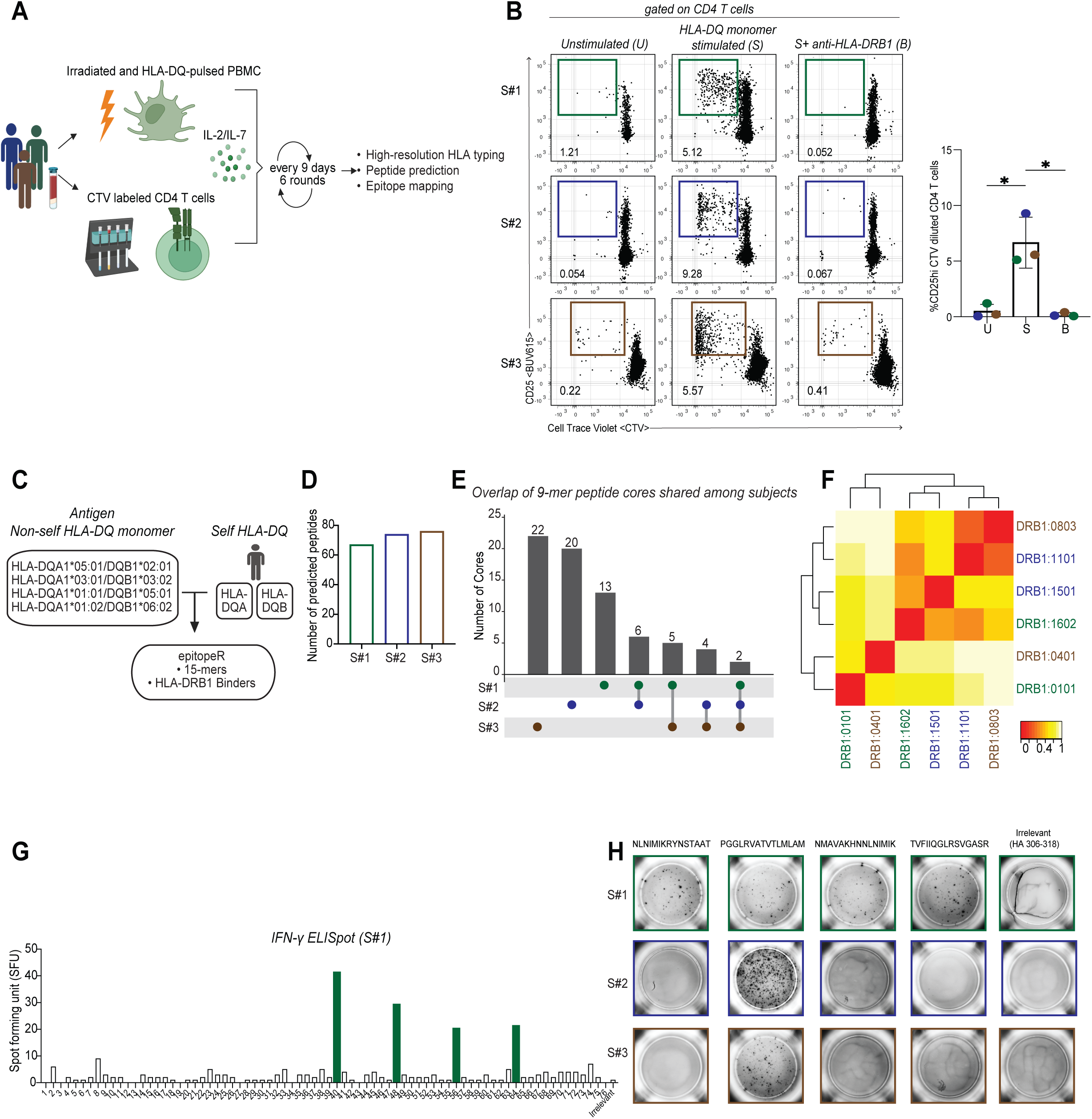
Identification of HLA-DQ-specific CD4 T cell responses in naïve individuals. (A) Schematic representation of the experimental strategy to detect HLA-DQ-specific CD4 T cells following enrichment with HLA-DQ monomers (n=3). (B) Representative flow cytometry plots showing CD25-expressing CTV-diluted CD4 T cells after the second round of stimulation with HLA-DQ-specific monomers (S), no stimulation (unstimulated, U), or stimulation with HLA-DQ-specific monomers in the presence of an anti-HLA-DRB1 antibody (B). The summary plot shows the percentage of CD25hi CTV-diluted CD4 T cells, with each subject represented by a distinct color. Data are presented as median ± s.d. and analyzed using the Mann-Whitney test (*p<0.05, ns = not significant). (C) Prediction pipeline for identifying non-self HLA-DQ-specific, HLA-DRB1-binding 15-mer MHC class II peptides using sequences from HLA-DQ monomers, analyzed with the epitopeR package. (D) Summary plot displaying the number of HLA-DQ-specific predicted peptides, with each subject represented by a distinct color. (E) UpSet plot illustrating the overlap of 9-mer HLA-DQ-specific peptide cores. Each dot, represented by a distinct color, corresponds to a single subject, and lines indicate overlaps. The y-axis denotes the number of shared 9-mer peptide cores. (F) Heatmap showing the similarity of MHC binding specific to the HLA-DRB1 alleles of subjects, with each subject uniquely represented by a distinct color. The scale represents the distance of similarity between presenting HLA alleles, where red (0) indicates high similarity and yellow (1) indicates low similarity. (G) Summary plot of epitope mapping for HLA-DQ-specific CD4 T cell epitopes using IFN-γ ELISpot, with results expressed as spot-forming units (SFU). (H) Representative IFN-γ ELISpot plots showing CD4 T cell responses to four immunogenic HLA-DQ-specific peptides and irrelevant influenza HA 306-318 peptide, with each subject uniquely represented by a distinct colo

To identify HLA-DQ specific CD4 T cell epitopes associated with HLA-DQ2.5, DQ5, DQ6, and DQ8, we used the epitope prediction software epitopeR to generate candidate 15-mer peptides that differ from the subjects’ self-HLA-DQ sequences while possessing the potential to bind to their HLA-DRB1 presenting alleles(26). This approach allowed us to identify putative non-self HLA-DQ-specific CD4 T cell epitopes that could stimulate an immune response (**Figure 1C, Supplemental Table 2**). The number of predicted peptides varied among subjects, with each subject presenting between 60 and 80 unique peptides (**Figure 1D**). We cross-referenced the predicted presentable peptides for each subject and identified 17 9-mer peptide cores that were shared by at least two subjects, indicating potential common immunogenic epitope cores or structural similarities between presenting HLA-DRB1 alleles between subjects (**Figure 1E**).

To assess the potential similarities of the binding motifs of our subjects’ HLA-DRB1 alleles, we employed the *in silico* tool MHCcluster to cluster the HLA-DRB1 alleles from the study subjects according to their predicted binding characteristics (27). Several alleles, including *DRB1*08:03*, *DRB1*11:01*, and *DRB1*16:02,* clustered closely together, suggesting that they may share structural features that underlie their similar binding specificities thereby contributing to the observed overlap of the predicted HLA-DQ peptides across subjects (**Figure 1F**). Following the peptide prediction, we synthesized candidate peptides and conducted functional assay to confirm their ability to elicit HLA-DQ-specific CD4 T cell responses using interferon-gamma (IFN-γ) enzyme-linked immunosorbent spot (ELISpot). The results, measured in spot-forming units (SFU), showed that four peptides induced elevated CD4 T cell responses compared to the baseline response elicited by an unrelated control peptide, HA306-318 (**Figures 1G and 1H**). Together, these findings demonstrate the effectiveness of *in silico* prediction methods, such as epitopeR, at narrowing a field of candidate HLA-DQ CD4 T cell epitopes in naïve individuals for further functional validation.

### 3.2. Enrichment with HLA-DQ monomers clonally expands HLA-DQ-reactive CD4 T cell clones with restricted TCRαβ usage

To examine the TCR repertoire of HLA-DQ-specific CD4 T cells and assess the impact of repeated stimulation with HLA-DQ monomers on TCR composition, we conducted TCR sequencing on samples obtained from a naïve individual at multiple time points, including pre-enrichment and post-enrichment rounds 4, 5, and 6 from the same subject (**Figure 2A**). Notably, the initial stimulation phases resulted in a decline in the cell count of expanded CD4 T cells, followed by a marked expansion beginning at post-enrichment round 4 (**Figure 2B**).

**Figure 2.**
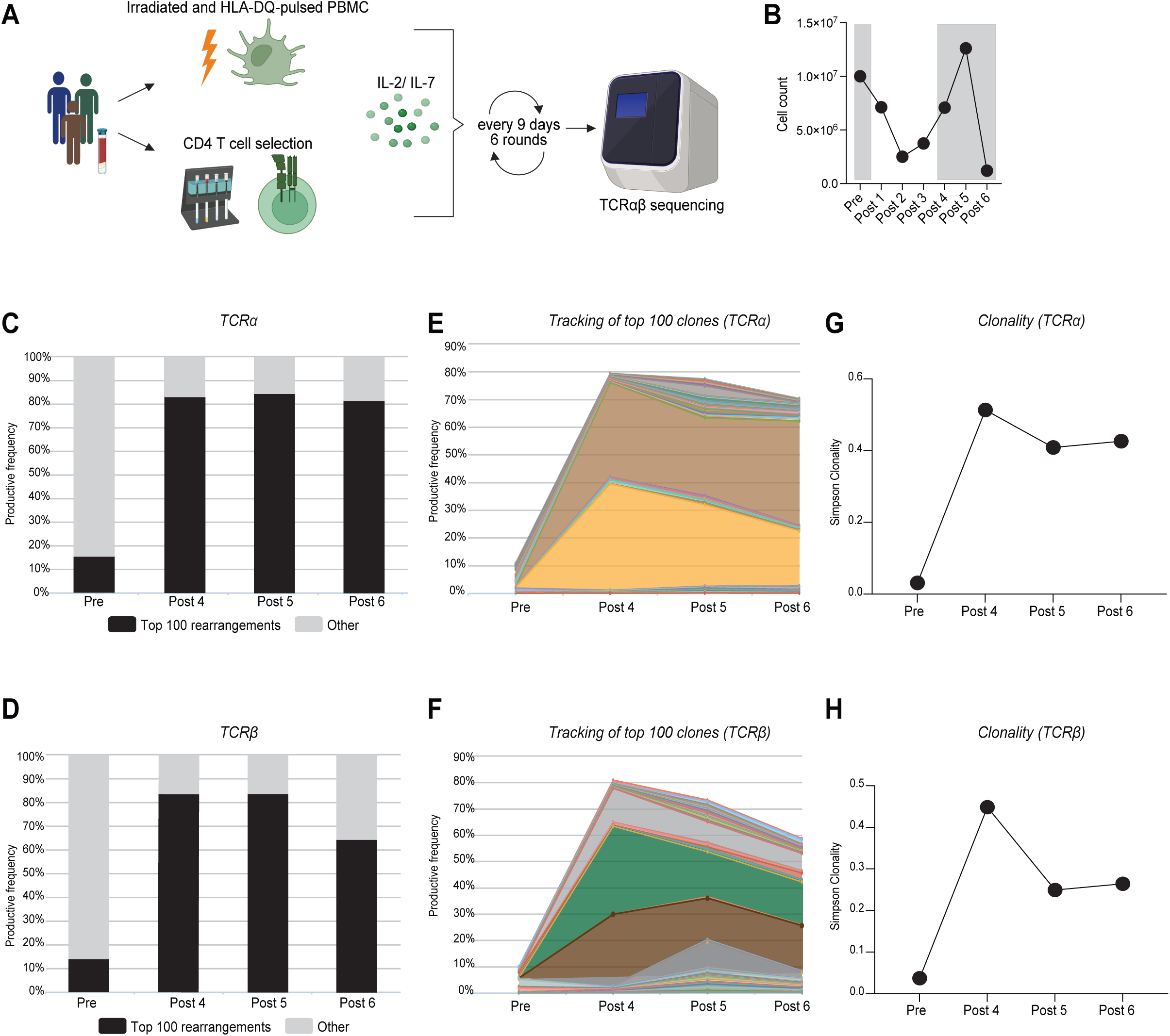
Clonal expansion of HLA-DQ-specific CD4 T cells following enrichment. (A) Schematic representation of the experimental strategy for unpaired TCRα and TCRβ sequencing of HLA- DQ-specific CD4 T cells at pre-enrichment and post-enrichment timepoints 4, 5, and 6. (B) Kinetics of total cell counts during T cell enrichment using HLA-DQ monomers. (C-D) Productive frequency of the top 100 TCRα (C) and TCRβ (D) rearrangements at pre-enrichment and post-enrichment timepoints 4, 5, and 6. (E-F) Longitudinal tracking of the top 100 TCRα (E) and TCRβ (F) rearrangements at pre-enrichment and post-enrichment timepoints 4, 5, and 6, with unique colors representing individual clonotypes. (G-H) Simpson clonality index of the TCRα (G) and TCRβ (H) repertoire at pre-enrichment and post-enrichment timepoints 4, 5, and 6.

TCR sequencing analysis revealed that the top 100 unique rearrangements of both TCRα and TCRβ in the post-enrichment samples (rounds 4, 5, and 6) accounted for 60-80% of the total TCR frequency, a significant increase compared to only 15% in the pre-enrichment sample (**Figures 2C and 2D**). Further tracking of the top 100 TCRα and TCRβ clonotypes revealed two dominant TCRα and three dominant TCRβ clonotypes that consistently expanded across the enrichment rounds (**Figures 2E and 2F**). The productive Simpson clonality index of HLA-DQ-specific CD4 T cells, which ranges from 0 to 1 (with values closer to 1 indicating a near-monoclonal population) (28), reflected these findings with a significant increase in post-enrichment samples (Figures 2G and 2H).

To identify which specific TCR clonotypes expanded over time, we conducted a further analysis of the TCR repertoire at each enrichment stage. In the pre-enrichment stage, the TCR repertoire was characterized by a diversity of small, polyclonal clonotypes, with the most prevalent TCRα and TCRβ clonotypes representing only 6% and 10% of the repertoire, respectively (**Figures 3A and 3B**). However, by post-enrichment round 4, a notable shift was observed, with two TCRα clonotypes and three TCRβ clonotypes significantly expanding. These dominant TCRα clonotypes now comprised 38% and 34% of the repertoire, while the TCRβ clonotypes constituted 33%, 26%, and 12% of the total (**Figures 3C and 3D**). Specifically, TCRAV21 and TCRAV26 emerged as the dominant TCRα genes, whereas TRBV18, TRBV27, and TRBV03 were the dominant TCRβ genes (**Figures 3C and 3D**). This pattern of expansion was consistently observed in the post-enrichment rounds 5 and 6, indicating that these specific clonotypes remained prominent throughout the T cell expansion process induced by HLA-DQ monomer stimulation (**Figures 3E-3H**). These findings suggest that repeated stimulation with HLA-DQ monomers effectively enriches a specific and oligoclonal TCR repertoire within CD4 T cells, characterized by the consistent expansion of select clonotypes.

**Figure 3.**
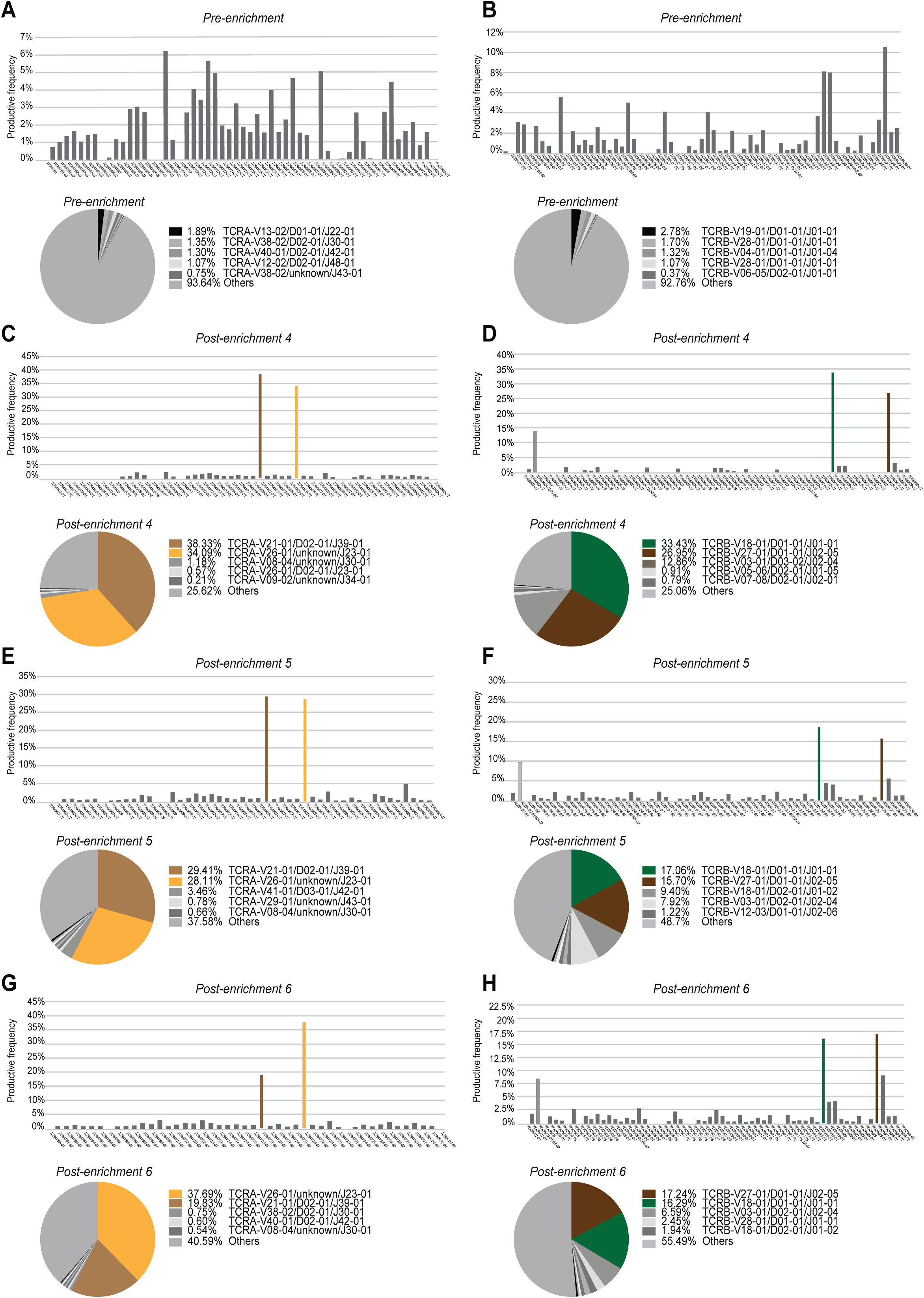
Restricted TCRαβ usage in clonally expanded HLA-DQ-specific CD4 T cells. (A-B) Productive frequency of TCRα (A) and TCRβ (B) gene usage in the pre-enrichment sample. The accompanying pie charts display the relative abundance of specific TCRα and TCRβ clonotypes. (C-D) Productive frequency of TCRα (C) and TCRβ (D) gene usage in the post-enrichment sample 4. The pie charts highlight the gene usage of two clonally expanded TCRα clonotypes (C) and three TCRβ clonotypes (D). (E-F) Productive frequency of TCRα (E) and TCRβ (F) gene usage in the post-enrichment sample 5. The pie charts highlight the gene usage of two clonally expanded TCRα clonotypes (E) and three TCRβ clonotypes (F). (G-H) Productive frequency of TCRα (G) and TCRβ (H) gene usage in the post-enrichment sample 6. The pie charts highlight the gene usage of two clonally expanded TCRα clonotypes (G) and three TCRβ clonotypes (H).

### 3.3. Tracking HLA-DQ-specific CD4 T cell clonotypes within the naïve TCR repertoire

To track HLA-DQ-specific CD4 T cell clonotypes in a naïve individual, we employed TCR sequencing to monitor clonotype frequencies across different stages of enrichment (**Figure 4A**). Samples were collected at three distinct stages: from the naïve T cell repertoire, pre-enrichment, and following repeated rounds of enrichment using HLA-DQ monomers, specifically after the fourth round of enrichment. The results demonstrated progressive enrichment of specific TCRα and TCRβ clonotypes over time, with a significant increase in their frequencies post-enrichment compared to pre-enrichment and the naïve repertoire (**Figure 4B and 4C**).

**Figure 4.**
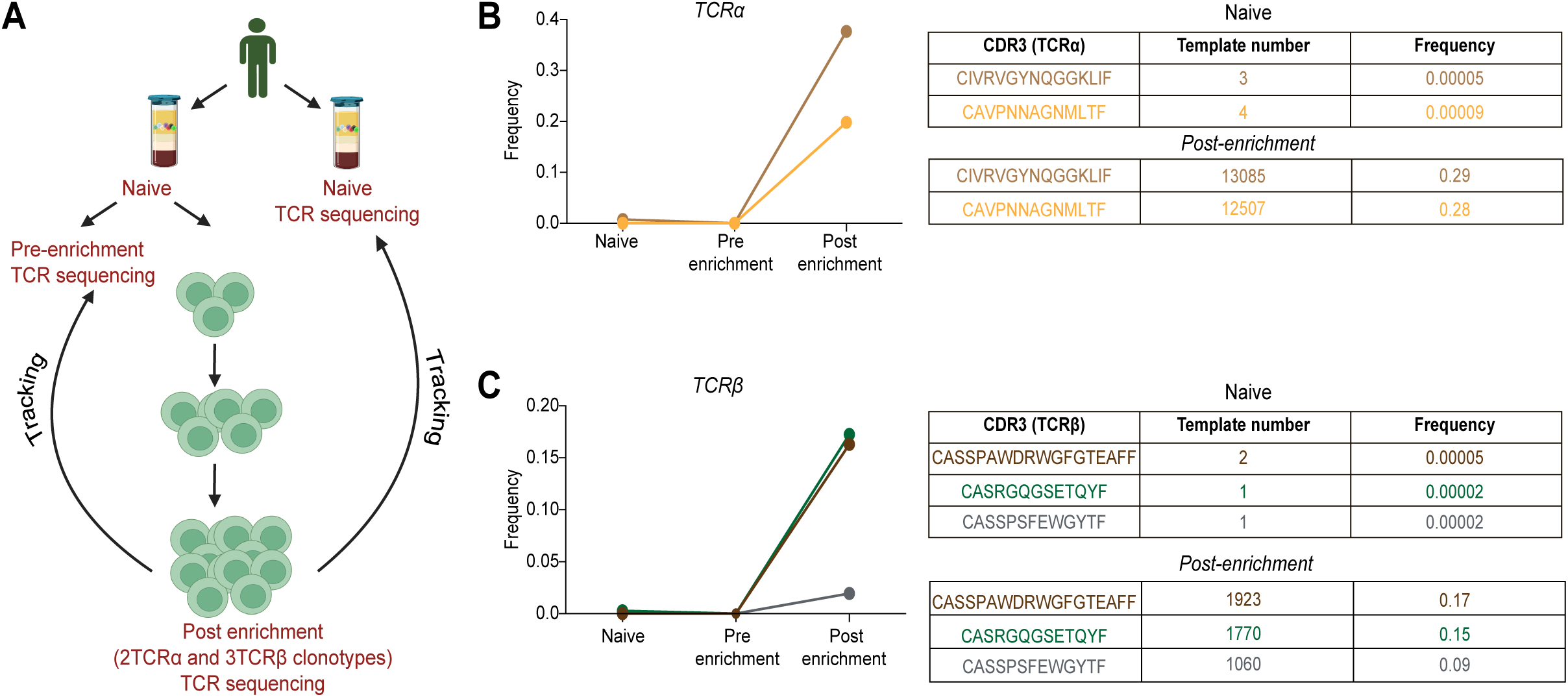
Tracking HLA-DQ-specific CD4 T cell clonotypes enriched from a naïve individual. (A) Schematic representation of the experimental strategy to track HLA-DQ-specific clonally expanded TCRα (2 clonotypes) and TCRβ (3 clonotypes). These clonotypes were identified through TCR sequencing at post-enrichment timepoint 4 and analyzed within the TCR repertoire at the pre-enrichment timepoint and the naïve repertoire from an independent sample of the same individual. (B) Frequency of the 2 clonally expanded HLA-DQ-specific TCRα clonotypes in the naïve, pre-enrichment, and post-enrichment TCR repertoire at timepoint 4, with each clonotype represented by a unique color. The corresponding table displays the template number and frequency of the clonally expanded TCRα clonotypes. (C) Frequency of the 3 clonally expanded HLA-DQ-specific TCRβ clonotypes in the naïve, pre-enrichment, and post-enrichment TCR repertoire at timepoint 4, with each clonotype represented by a unique color. The corresponding table displays the template number and frequency of the clonally expanded TCRβ clonotypes.

Two clonally expanded TCRα clonotypes observed at the post-enrichment stage were tracked back to both the pre-enrichment stage and the naïve TCR repertoire from the same individual. Specifically, the template count of the TCRα clonotype *CIVRVGYNQGGKLIF* increased from 3 in the naïve repertoire to 13,085 post-enrichment, and *CAVPNNAGNMLTF* increased from 4 to 12,507 (**Figure 4B**). Correspondingly, their frequencies rose from 0.0005 to 0.29 and 0.0009 to 0.28, respectively. Similarly, the template count of the TCRβ clonotype *CASSPAWDRWGFGTEAFF* increased from 2 in the naïve repertoire to 1,923 post-enrichment (**Figure 4C**). Additionally, clonotypes *CASRGQGSETQYF* and *CASSPSFEWGYTF*, which initially had template counts of 1 each, increased to 1,770 and 1,060, respectively. Their frequencies rose from 0.00005 to 0.17 for *CASSPAWDRWGFGTEAFF*, from 0.00002 to 0.15 for *CASRGQGSETQYF*, and from 0.00002 to 0.09 for *CASSPSFEWGYTF*. Together, these findings highlight the capability of the enrichment process to selectively amplify rare HLA-DQ-specific T cell clonotypes from the naïve repertoire.

## 4. Discussion

Understanding donor-specific T cell responses in naive individuals is essential for advancing transplant immunology, as it provides insights into immune recognition of donor antigens without prior sensitization. Naive CD4 T cells, characterized by high CD62L and CCR7 expression, continuously circulate in the periphery between secondary lymphoid organs, positioning them as primary drivers of initial allograft rejection (29). Their unprimed, polyclonal, and diverse TCR repertoire enables recognition of a broad range of donor antigens, challenging long- term graft survival in unsensitized recipients (29). Thus, defining the donor-specific T cell repertoire in naive individuals is critical for identifying precursor cells with rejection potential and predicting donor-specific immune responses prior to transplantation.

In transplantation, HLA-DQ DSA is increasingly recognized as a key factor in chronic allograft rejection, with significant implications for graft survival (4–7). While the clinical impacts of DSA targeting HLA-A, -B, and -DR loci are well documented, HLA-DQ-directed DSA remains comparatively underexplored (23). Given its clinical importance, this study investigates HLA-DQ-specific CD4 T cell responses in naive individuals, demonstrating that HLA-DQ monomers can serve as effective alloantigen sources. This approach enhances sensitivity in detecting HLA-DQ-specific CD4 T cells, facilitates the identification of HLA-DQ epitopes, and supports characterization of the corresponding TCR repertoire in naïve individuals, while also offering the potential to be extended to other HLA monomers for similar applications and genetic expression of whole HLA proteins.

One of the key challenges in transplant immunology, particularly in humans, is the paucity of reliable tools to detect donor-specific T cells (30,31). Existing techniques struggle to identify the small populations of donor- specific alloreactive T cells, making their early detection difficult. To address this limitation, we developed an assay to detect HLA-DQ-specific CD4 T cell responses in naive individuals using HLA-DQ monomers representing the prevalent HLA-DQ2.5, DQ5, DQ6, and DQ8 antigens (23). An advantage of using HLA monomers as intact donor MHC is that T cells stimulated in these assays may closely mimic the behavior of T cells in the polyclonal repertoire activated in the context of transplantation against engrafted tissue, as a result of the natural processing of donor antigens by antigen-presenting cells (APCs). By excluding self-HLA alleles from the stimulation cocktail, we ensured that the T cell responses observed were specific to alloantigens. CD4 T cell proliferation and activation marker upregulation validated the enrichment of HLA-DQ-specific T cells. These results highlight the potential utility of HLA monomers in monitoring alloantigen-specific responses in the context of human transplantation.

To identify specific HLA-DQ CD4 T cell epitopes, we employed the epitope prediction tool epitopeR, by selecting 15-mer peptides predicted to bind self-HLA-DRB1 alleles. Unlike previous studies relying on overlapping peptide panels (32,33), our *in silico* approach allowed high-throughput selection of peptides with strong binding affinity and shared 9-mer cores across multiple individuals. MHC specificity heatmaps revealed clusters of donor-specific HLA-DRB1 alleles with similar binding properties, notably *DRB1*\**08:03, DRB1*11:01*, and *DRB1*\**16:02*, suggesting consistent antigen presentation patterns that may influence CD4 T cell activation profiles (34). IFN-γ ELISpot assays functionally validated several HLA-DQ-specific peptides, which elicited responses across individuals, likely due to conserved antigen presentation features among HLA-DRB1 alleles.

Further tracking of the TCR repertoire in HLA-DQ-specific CD4 T cells revealed a shift toward oligoclonality with repeated HLA-DQ monomer stimulations, suggesting the preferential expansion of specific TCR clonotypes. TCRαβ sequencing corroborated this by showing an increased prevalence of particular TCRα and TCRβ clonotypes over time, with dominant clonotypes emerging after repeated stimulation. Notably, these clonotypes were traceable back to the naïve TCR repertoire, demonstrating that rare precursor T cells with donor-specific reactivity can be selectively enriched through iterative stimulation. This selective enrichment reflects the activation of high-affinity TCRs specific to HLA-DQ-restricted donor antigens and is consistent with prior studies documenting restricted TCR gene usage among alloreactive T cells (35). Tracking these clonally expanded TCRs underscores their potential as one of the key drivers of alloreactive immune responses and suggests that targeting these clonotypes could potentially suppress or prevent alloimmune rejection.

Together, this study highlights the utility of HLA-DQ monomers as potential tools for detecting and characterizing donor-specific CD4 T cell responses in naive individuals. By combining peptide prediction, functional validation, and TCR repertoire analysis, our findings provide insights into the immunogenicity of HLA-DQ molecules. Expanding this approach to encompass broader HLA contexts holds promise for advancing personalized transplant strategies and improving patient outcomes.

## Limitations

While our findings highlight the importance of HLA-DRB1 in CD4 T cell activation under these experimental conditions, it is worth noting that HLA-DQ or HLA-DP molecules may also present donor-derived peptides in other contexts. The potential role of these molecules, particularly in diverse antigen-processing scenarios or transplantation settings, warrants further investigation. These possibilities underscore the broader complexity of donor-specific immune responses and the need for continued exploration of all HLA class II molecules.

## Acknowledgements

This work was financially supported with funds from the *Trailsend fund for Immunotherapy* gift from the Trailsend Foundation directed by James C. Kennedy (C.P.L.); National Institutes of Health grants 5U01AI138909-05 (C.P.L.); Carlos and Marguerite Mason Trust sponsored *Mason Trust Research grants* (C.P.L.); CUREIT: Curing the Uncurable via RNA-Encoded Immunogene Tuning; Sponsor: ARPA-H; 1AY1AX000001-01 (C.P.L. and H.T.K.); Cancer Research Institute Lloyd J. Old STAR program (H.T.K.)

We want to acknowledge J.D. Altman (Emory University) for assistance in conceptualizing the immunopeptidomics. Finally, we would like to acknowledge all the members of the Larsen and Kissick Lab for their helpful contributions and insightful discussions regarding this work.

## Author Contributions

Z.Z., H.T.K., and C.P.L. designed the study and analyzed data. Z.Z. and D.C. wrote the manuscript. Z.Z. performed the experiments. A.H. processed the blood samples. Z.Z., X.Z., and C.P.L. performed epitope prediction. H.T.K. and C.P.L. supervised the research. All the authors reviewed the manuscript.

## Data and Software Availability

Custom Python and R scripts for Epitope Prediction and TCR analysis are available upon request.

## Supporting information statement

Additional supporting information may be found online in the Supporting Information Section.

## Reference

1. Colvin RB, Smith RN. Antibody-mediated organ-allograft rejection. Nat Rev Immunol (2005) 5:807–817. doi: 10.1038/nri1702

2. Lachmann N, Terasaki PI, Budde K, Liefeldt L, Kahl A, Reinke P, Pratschke J, Rudolph B, Schmidt D, Salama A, et al. Anti-Human Leukocyte Antigen and Donor-Specific Antibodies Detected by Luminex Posttransplant Serve as Biomarkers for Chronic Rejection of Renal Allografts. Transplantation (2009) 87:1505–1513. doi: 10.1097/TP.0b013e3181a44206

3. Everly MJ, Rebellato LM, Haisch CE, Ozawa M, Parker K, Briley KP, Catrou PG, Bolin P, Kendrick WT, Kendrick SA, et al. Incidence and Impact of De Novo Donor-Specific Alloantibody in Primary Renal Allografts. Transplantation (2013) 95:410–417. doi: 10.1097/TP.0b013e31827d62e3

4. Willicombe M, Brookes P, Sergeant R, Santos-Nunez E, Steggar C, Galliford J, McLean A, Cook TH, Cairns T, Roufosse C, et al. De novo DQ donor-specific antibodies are associated with a significant risk of antibody-mediated rejection and transplant glomerulopathy. Transplantation (2012) 94:172–177. doi: 10.1097/TP.0b013e3182543950

5. Zhang X, Kransdorf E, Levine R, Patel JK, Kobashigawa JA. HLA-DQ mismatches stimulate de novo donor specific antibodies in heart transplant recipients. Hum Immunol (2020) 81:330–336. doi: 10.1016/j.humimm.2020.04.003

6. Zhang X, Reinsmoen NL, Kobashigawa JA. HLA Mismatches Identified by a Novel Algorithm Predict Risk of Antibody-mediated Rejection From De Novo Donor-specific Antibodies. Transplantation (2024) doi: 10.1097/TP.0000000000005140

7. Ozawa M, Rebellato LM, Terasaki PI, Tong A, Briley KP, Catrou P, Haisch CE. Longitudinal testing of 266 renal allograft patients for HLA and MICA antibodies: Greenville experience. Clin Transpl (2006)265–290.

8. Zhanzak Z, Cina D, Johnson A, Larsen CP. Implications of MHC-restricted Immunopeptidome in Transplantation. (2024) 15: doi: 10.3389/fimmu.2024.1436233

9. Geiger R, Duhen T, Lanzavecchia A, Sallusto F. Human naive and memory CD4+ T cell repertoires specific for naturally processed antigens analyzed using libraries of amplified T cells. J Exp Med (2009) 206:1525–1534. doi: 10.1084/jem.20090504

10. Stone JD, Chervin AS, Kranz DM. T-cell receptor binding affinities and kinetics: impact on T-cell activity and specificity. Immunology (2009) 126:165. doi: 10.1111/j.1365-2567.2008.03015.x

11. Joglekar AV, Li G. T cell antigen discovery. Nat Methods (2021) 18:873–880. doi: 10.1038/s41592-020-0867-z

12. Christophersen A. Peptide-MHC class I and class II tetramers: From flow to mass cytometry. HLA (2020) 95:169–178. doi: 10.1111/tan.13789

13. Bentzen AK, Marquard AM, Lyngaa R, Saini SK, Ramskov S, Donia M, Such L, Furness AJS, McGranahan N, Rosenthal R, et al. Large-scale detection of antigen-specific T cells using peptide-MHC-I multimers labeled with DNA barcodes. Nat Biotechnol (2016) 34:1037–1045. doi: 10.1038/nbt.3662

14. Zhang S-Q, Ma K-Y, Schonnesen AA, Zhang M, He C, Sun E, Williams CM, Jia W, Jiang N. High-throughput determination of the antigen specificities of T cell receptors in single cells. Nat Biotechnol (2018) 36:1156–1159. doi: 10.1038/nbt.4282

15. Ng AHC, Peng S, Xu AM, Noh WJ, Guo K, Bethune MT, Chour W, Choi J, Yang S, Baltimore D, et al. MATE-Seq: microfluidic antigen-TCR engagement sequencing. Lab Chip (2019) 19:3011–3021. doi: 10.1039/C9LC00538B

16. Lee RS, Yamada K, Houser SL, Womer KL, Maloney ME, Rose HS, Sayegh MH, Madsen JC. Indirect recognition of allopeptides promotes the development of cardiac allograft vasculopathy. Proc Natl Acad Sci (2001) 98:3276–3281. doi: 10.1073/pnas.051584498

17. Lee RS, Grusby MJ, Glimcher LH, Winn HJ, Auchincloss H. Indirect recognition by helper cells can induce donor-specific cytotoxic T lymphocytes in vivo. J Exp Med (1994) 179:865–872. doi: 10.1084/jem.179.3.865

18. Vella JP, Magee C, Vos L, Womer K, Rennke H, Carpenter CB, Hancock W, Sayegh MH. Cellular and humoral mechanisms of vascularized allograft rejection induced by indirect recognition of donor MHC allopeptides. Transplantation (1999) 67:1523–1532. doi: 10.1097/00007890-199906270-00005

19. Glanville J, Huang H, Nau A, Hatton O, Wagar LE, Rubelt F, Ji X, Han A, Krams SM, Pettus C, et al. Identifying specificity groups in the T cell receptor repertoire. Nature (2017) 547:94–98. doi: 10.1038/nature22976

20. Dash P, Fiore-Gartland AJ, Hertz T, Wang GC, Sharma S, Souquette A, Crawford JC, Clemens EB, Nguyen THO, Kedzierska K, et al. Quantifiable predictive features define epitope-specific T cell receptor repertoires. Nature (2017) 547:89–93. doi: 10.1038/nature22383

21. Zhang W, Hawkins PG, He J, Gupta NT, Liu J, Choonoo G, Jeong SW, Chen CR, Dhanik A, Dillon M, et al. A framework for highly multiplexed dextramer mapping and prediction of T cell receptor sequences to antigen specificity. Sci Adv (2021) 7:eabf5835. doi: 10.1126/sciadv.abf5835

22. Lefranc M-P, Giudicelli V, Ginestoux C, Jabado-Michaloud J, Folch G, Bellahcene F, Wu Y, Gemrot E, Brochet X, Lane J, et al. IMGT(R), the international ImMunoGeneTics information system(R). Nucleic Acids Res (2009) 37:D1006–D1012. doi: 10.1093/nar/gkn838

23. Lee H, Min JW, Kim J-I, Moon I-S, Park K-H, Yang CW, Chung BH, Oh E-J. Clinical Significance of HLA-DQ Antibodies in the Development of Chronic Antibody-Mediated Rejection and Allograft Failure in Kidney Transplant Recipients. Medicine (Baltimore) (2016) 95:e3094. doi: 10.1097/MD.0000000000003094

24. Breman E, Van Miert PP, Van Der Steen DM, Heemskerk MH, Doxiadis II, Roelen D, Claas FH, Van Kooten C. HLA Monomers as a Tool to Monitor Indirect Allorecognition. Transplantation (2014) 97:1119–1127. doi: 10.1097/TP.0000000000000113

25. Wölfl M, Greenberg PD. Antigen-specific activation and cytokine-facilitated expansion of naive, human CD8+ T cells. Nat Protoc (2014) 9:950–966. doi: 10.1038/nprot.2014.064

26. Zhanzak Z, Johnson AC, Foster P, Cardenas MA, Morris AB, Zhang J, Karadkhele G, Badell IR, Morris AA, Au-Yeung BB, et al. Identification of indirect CD4+ T cell epitopes associated with transplant rejection provides a target for donor-specific tolerance induction. Immunity (2025)S1074761325000330. doi: 10.1016/j.immuni.2025.01.008

27. Thomsen M, Lundegaard C, Buus S, Lund O, Nielsen M. MHCcluster, a method for functional clustering of MHC molecules. Immunogenetics (2013) 65:655–665. doi: 10.1007/s00251-013-0714-9

28. Chiffelle J, Genolet R, Perez MA, Coukos G, Zoete V, Harari A. T-cell repertoire analysis and metrics of diversity and clonality. Curr Opin Biotechnol (2020) 65:284–295. doi: 10.1016/j.copbio.2020.07.010

29. Golshayan D, Wyss J-C, Buckland M, Hernandez-Fuentes M, Lechler RI. Differential role of naïve and memory CD4 T-cell subsets in primary alloresponses. Am J Transplant Off J Am Soc Transplant Am Soc Transpl Surg (2010) 10:1749–1759. doi: 10.1111/j.1600-6143.2010.03180.x

30. Young JS, McIntosh C, Alegre M-L, Chong AS. Evolving Approaches in the Identification of Allograft- Reactive T and B Cells in Mice and Humans. Transplantation (2017) 101:2671–2681. doi: 10.1097/TP.0000000000001847

31. Jiang X, Shi QS, Wu C-Y, Xu L, Yang H, Medhat Askar. Investigative and laboratory assays for allogeneic rejection – A clinical perspective. Transplant Rep (2023) 8:100133. doi: 10.1016/j.tpr.2023.100133

32. Vella JP, Spadafora-Ferreira M, Murphy B, Alexander SI, Harmon W, Carpenter CB, Sayegh MH. Indirect allorecognition of major histocompatibility complex allopeptides in human renal transplant recipients with chronic graft dysfunction. Transplantation (1997) 64:795–800. doi: 10.1097/00007890-199709270-00001

33. Reznik SI, Jaramillo A, SivaSai KSR, Womer KL, Sayegh MH, Trulock EP, Patterson GA, Mohanakumar T. Indirect Allorecognition of Mismatched Donor HLA Class II Peptides in Lung Transplant Recipients with Bronchiolitis Obliterans Syndrome. Am J Transplant (2001) 1:228–235. doi: 10.1034/j.1600-6143.2001.001003228.x

34. Álvaro-Benito M, Abualrous ET, Lingel H, Meltendorf S, Holzapfel J, Sticht J, Kuropka B, Clementi C, Kuppler F, Brunner-Weinzierl MC, et al. Effective assessment of CD4^+^ T cell Immunodominance patterns: impact of antigen processing and HLA restriction. (2024) doi: 10.1101/2024.01.10.574975

35. Liu Z, Sun YK, Xi YP, Hong B, Harris PE, Reed EF, Suciu-Foca N. Limited usage of T cell receptor V beta genes by allopeptide-specific T cells. J Immunol Baltim Md 1950 (1993) 150:3180–3186.

